# Low intrinsic efficacy alone cannot explain the improved side effect profiles of new opioid agonists

**DOI:** 10.1101/2020.11.19.390518

**Authors:** Edward L. Stahl, Laura M. Bohn

**Affiliations:** Department of Molecular Medicine, The Scripps Research Institute, 130 Scripps Way, Jupiter, FL, 33458

## Abstract

In a recent report in Science Signaling (DOI: 10.1126/scisignal.aaz3140), it was suggested that low intrinsic agonism, and not biased agonism, leads to an improvement in the separation of potency in opioid-induced respiratory suppression versus antinociception. Although many of the compounds that were tested have been shown to display G protein signaling bias in prior publications, the authors conclude that since they cannot detect biased agonism in their cellular signaling studies the compounds are therefore not biased agonists. Rather, they conclude that it is low intrinsic efficacy that leads to the therapeutic window improvement. Intrinsic efficacy is the extent to which an agonist can stimulate a G protein-coupled receptor (GPCR) response in a system, while biased agonism takes into consideration not only intrinsic efficacy, but also potency of an agonist in an assay. Herein, we have re-analyzed the data presented in the published work (DOI: 10.1126/scisignal.aaz3140) (including the recent Erratum: DOI: 10.1126/scisignal.abf9803) to derive intrinsic efficacy and bias factors as ΔΔlog(τ/K_A_) and ΔΔlog(Emax/EC_50_). Based on this reanalysis, the data support the conclusion that biased agonism, favoring G protein signaling, was observed. Moreover, a conservation of rank order intrinsic efficacy was not observed upon comparing responses in each assay, further suggesting that multiple active receptor states were present. These observations agree with prior studies wherein oliceridine, PZM21 and SR-17018 were first described as biased agonists with improvement in antinociception over respiratory suppression in mice. Therefore, the data in the Science Signaling manuscript does provide strong corroborating evidence that G protein signaling bias may be a means to improve opioid analgesia while avoiding certain undesirable side effects.

## Introduction

In the March 31, 2020 issue of Science Signaling, a research article by Gillis et al. ^*1, 2*^ (hereafter referred to as the SS-manuscript) investigated a series of mu opioid agonists for activity in a compendium of *in vitro* cell-based bioluminescence resonance energy transfer (BRET) studies (summarized in **Table 1**) to evaluate the pharmacological basis of functional selectivity. In particular, the authors focused on two discrete avenues of response: those medicated by G protein signaling or βarrestin2 recruitment. The SS-manuscript also included studies in mice that were intended to compare the therapeutic window of the selected compounds by determining potency in agonist-induced antinociception (hot-plate latency) and respiratory suppression (respiratory frequency) measures. These physiological responses are relevant as they reflect a similar relationship to the therapeutic window observed in clinical use of known opioids. While the study confirmed that the selected biased agonists showed an improved therapeutic window, the authors failed to detect biased agonism in their signaling assays and conclude that intrinsic efficacy and not biased agonism was responsible for the improved therapeutic window ^*1*^.

**Table 1.**
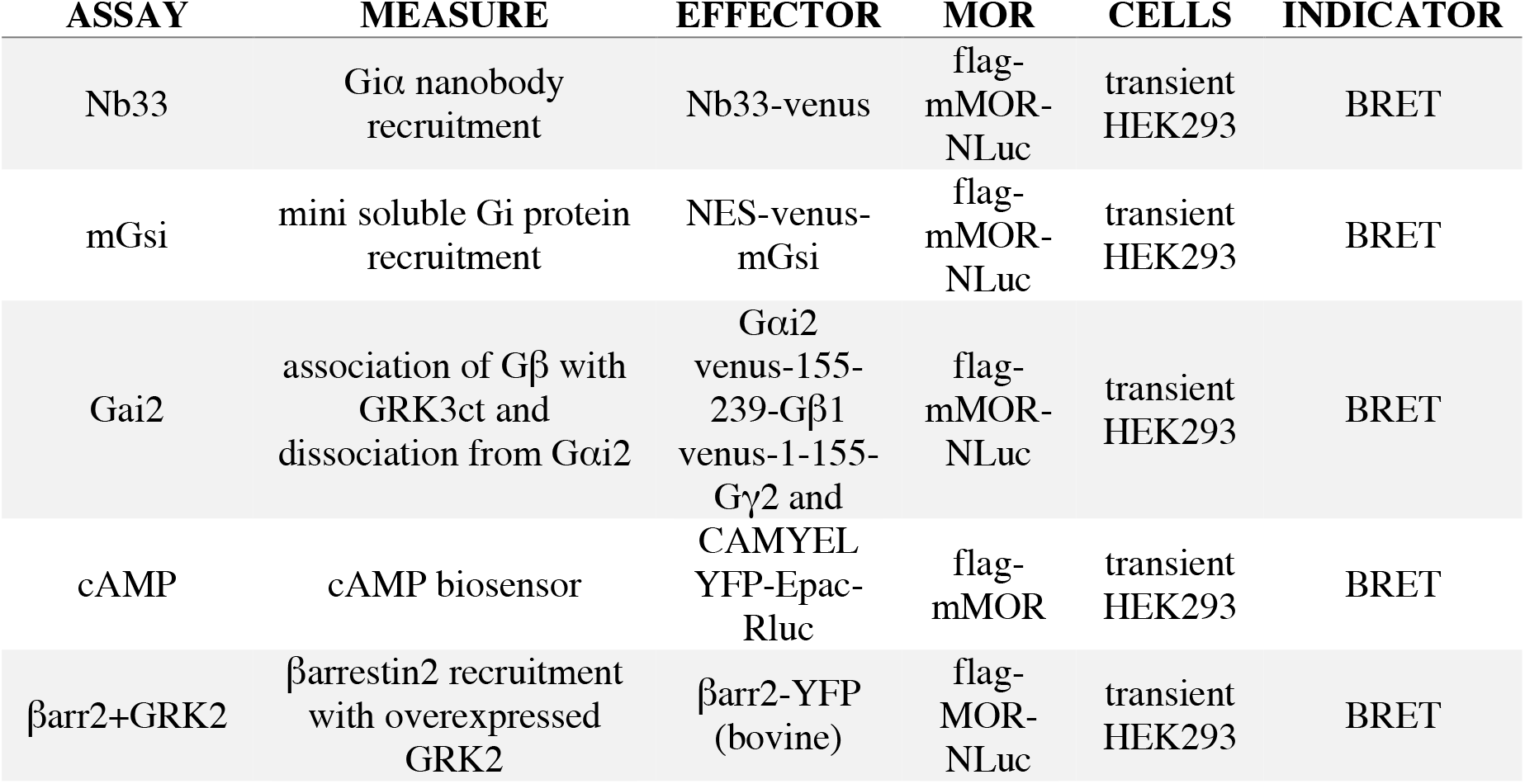
Assay abbreviations from the SS-manuscript ^*1*^.

The pursuit of agonists that display G protein signaling over βarrestin2 recruitment was inspired by early observations of mice lacking βarrestin2; where antinociception was enhanced and respiratory suppression was diminished in response to morphine ^*3-5*^. Several laboratories have developed such agonists, including PZM21 ^*6*^, a series of Scripps Research “ SR” compounds ^*7, 8*^, a fungal natural product ^*9*^, derivatives of mitragynine ^*10*^ and importantly, the first G protein signaling-biased MOR agonist, oliceridine (TRV-130, Olinvyk^®^,^*11*^) that has progressed to the clinic ^*12*^. Moreover, an improvement in producing antinociception with fewer signs of respiratory suppression has been seen for these compounds in rodents (with the exception of the fungal agonist which was not tested in vivo). Importantly, with the emerging clinical studies demonstrating an improvement in analgesic efficacy over respiratory suppression for oliceridine ^*13, 14*^, it would seem that the early mouse studies were predictive of a useful strategy for improving opioid therapeutics by developing G protein signaling-biased agonists.

Determining whether a compound shows a preference for signaling in one pathway over another has proven to be less straightforward than anticipated. These measures are complicated by the use of overexpression systems in different cellular backgrounds, varying receptor levels, tags and modifications to the receptors and effectors to amplify signal and the assumption that a plethora of assays designed to detect G protein mediated signaling are equal surrogates for that response. This leads to the assumption that the determination of potency (EC_50_) and efficacy (Emax) across a diverse array of “ G protein signaling” assays will define the agonist’s ability to stimulate G proteins. In reality, receptor number, signal amplification and the sensitivity of the assay will greatly affect the determination of potency and efficacy; however, if all agonists have an equivalent propensity to activate different pathways, then their rank order efficacy should be preserved (the most potent and efficacious agonist should remain so in all assays) ^*15-17*^.

In the SS-manuscript ^*1*^, it is proposed that ligand rank order intrinsic efficacy and *not* ligand bias is the driving factor behind the separation of antinociception from respiratory suppression of PZM21, oliceridine, buprenorphine and SR-17018. The authors quantified “ biased agonism” and agonist intrinsic efficacy across cellular signaling assays using the operational model originally described by Black and Leff ^*18*^. Notably, the authors reanalyzed efficacy (E_max_) values from several recent papers (including PZM21 ^*6*^ and SR-17018 ^*8*^ to support the proposal that it is intrinsic efficacy, and not biased agonism, that leads to improvements in the therapeutic window ^*1, 19, 20*^. *The conclusions of this report* ^*1*^ *and the value of partial agonist efficacy has been the subject of a number of recent review papers* ^*19-21*^.

The operational model has become common place in studying agonist performance across a variety of experimental systems and serves as a means to quantify biased agonism. The key feature that makes this model appealing is its capacity to produce a simplified correction for non-linear occupancy-response relationships. The model employs a hyperbolic occupancy-response relationship and assumes that, at a partial level of agonist occupancy, some agonists are able to fully activate their effector proteins. The product of this analysis are two unique parameters, tau (τ) and K_A_. K_A_ is an equilibrium affinity constant which describes the avidity by which an agonist-receptor complex is formed, while τ is an (intrinsic) efficacy parameter that describes how well the agonist-receptor complex is able to convey a signal to the effector protein. An easy way to understand the scale of τ is that when an agonist has an Emax of 50%, it will have a τ = 1 ^*22*^. This can be realized by substitution of τ = 1 into Equation 1, from Black and Leff ^*18*^: **Equation 1**.

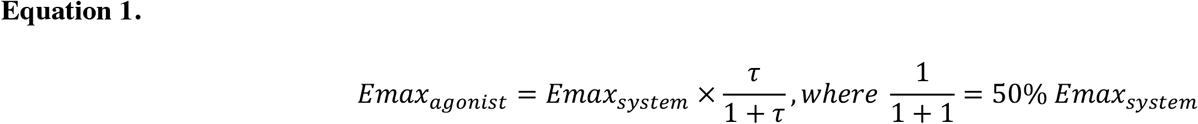

As the Emax of the agonist approaches 0, the value of tau will approach 0. For most experimental systems, however, τ will be a positive value as inverse agonist receptor reserve is not commonly encountered and would require a different set of fundamental assumptions about the system. Since τ is positive, it can be expressed as log(τ) which will then extend across positive and negative numbers.

In the SS-manuscript, the operational model is employed to calculate intrinsic efficacy (τ) for several partial agonists to assess MOR engagement with G protein or other effectors relative to the 100% maximum system response produced by DAMGO (see Table 1 for assay description, abbreviated names of the signaling assays will be used throughout the text). The resulting values for log(τ) (SS-Table 1), Emax (SS-Table 2), EC_50_ (SS-Table 3), and Δlog(τ/K_A_) (SS-Table S1) are presented in the SS-manuscript (for clarity, figures and tables in the original manuscript will be referred to with the prefix: “ SS-,” i.e. SS-Table 1 ^*1*^). An erratum ^*2*^, wherein the following changes were made: SS-Table S1 (Δlog(τ/K_A_)) values for cAMP for all compounds were replaced, and buprenorphine values were added for the βarr2 and βarr2 (+GRK2); SS-Table 3 (EC_50_) values for SR-17018 in βarr2 (+GRK2) were changed; no graphs or conclusions were changed in the erratum. All analyses presented herein include the “ erratum” values.

## Analysis

### Discrepancies between τ and Emax

Since the primary conclusion of the manuscript is that intrinsic efficacy, and not agonist bias, correlates with an improved therapeutic window, it was reasonable to investigate these measures of τ and Emax. As noted, the τ value can be calculated from Emax, and vice versa (Equation 1, ^*1, 18*^); therefore, Emax values presented after conversion from SS-Table 1 (conversion using substitution into Equation 1) were compared to the Emax values presented in SS-Table 2 in **Figure 1**. For this comparison, only partial agonists and only the assays that presented all of the values in both SS-Tables were included. While morphine shows the expected consistency between the Emax derived from the two tables; there is considerably less agreement for other agonists.

**Figure 1.**
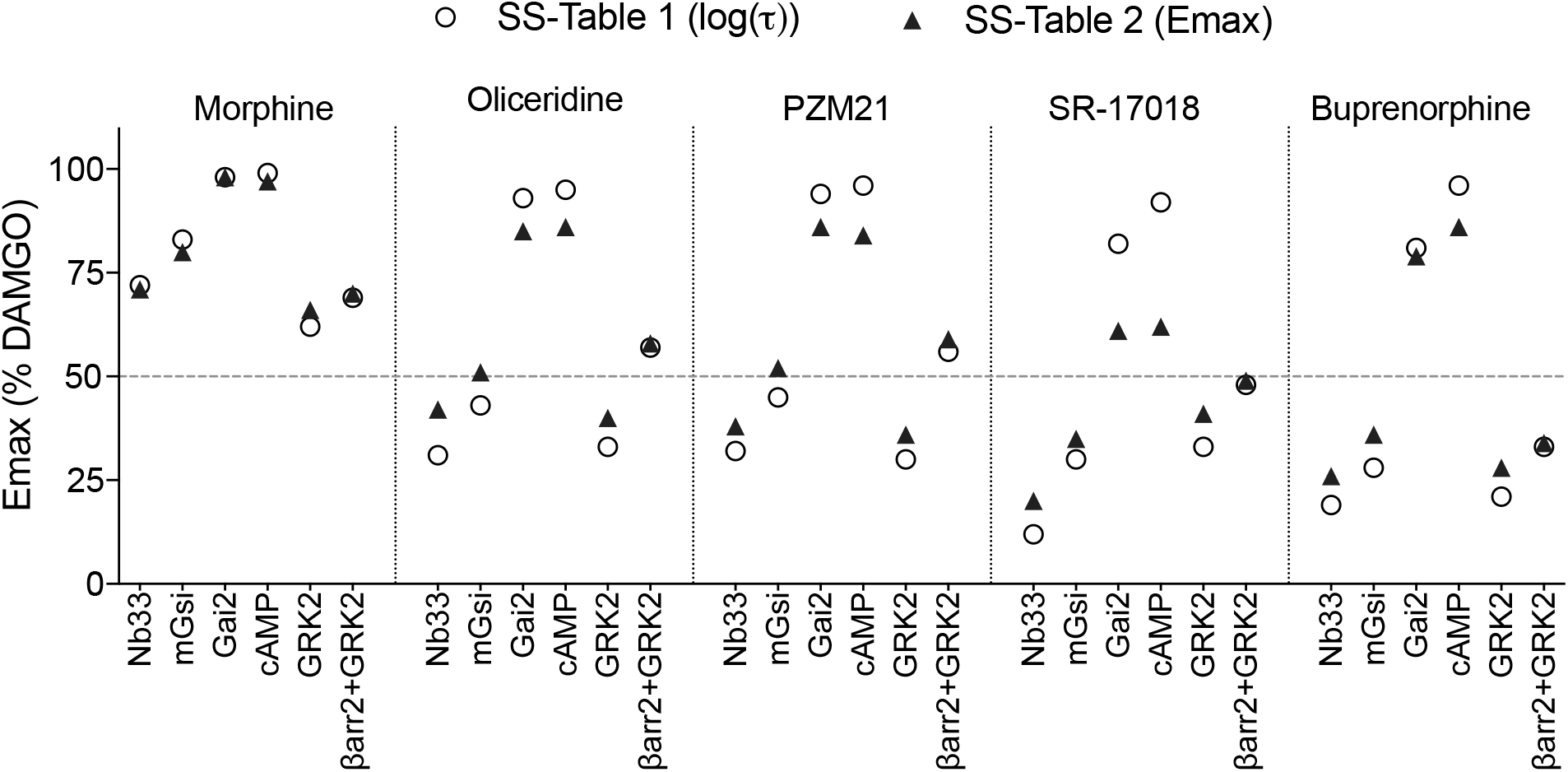
Emax is graphically presented from the values provided in SS-Table 2 in comparison to the Emax calculated from log(τ) (SS-Table 1) using Equation 1.

Interestingly, the Emax is overestimated when derived from the log(τ) and this is consistent for most of the amplified assays (Gαi2 and cAMP) while underestimated for the assays that most proximally detect G protein-receptor interaction (Nb33 and mGsi). The discrepancy is most pronounced for the data provided for SR-17018 where there is significant separation between the Emax prediction from τ (SS-Table 1) and the Emax estimate (SS-Table 2). Furthermore, it is remarkable that certain log(τ) values ± SEM in SS-Table 1 are negative. Specifically, while the Emax for oliceridine was reported as 51% for mGsi in SS-Table 2, a negative log(τ) value of -0.12 ± 0.05 was presented in SS-Table 1 which would require that the Emax be less than 50%, by definition. It should be mentioned that although the mean ± SEM does not overlap zero, this information is secondary to the primary Emax vs. log(τ) comparison. Better stated, a mean Emax > 50% cannot agree with a mean log(τ) < 0 regardless of any consideration of standard error of the mean. Overall, these discrepancies raise questions about the utility of the determination of τ, as presented in SS-Table 1, for assigning rank order intrinsic efficacy.

The use of the operational model centrally assumes a non-linear occupancy-response relationship. As such, an agonist response curve (potency) is expected to reside to the left of the agonist occupancy binding curve (affinity). That is, the agonist affinity is amplified by the system to produce the observed agonist EC_50_ in a given response. Concern is warranted when a hyperbolic occupancy-response function is employed to analyze what would be better described as a linear occupancy-response relationship i.e., when there is minimal separation between the response and binding curve ^*23, 20*^. Specifically, this can alter the way τ and K_A_ relate to Emax and EC_50_ due to an inappropriate correction for system amplification. However, as τ and K_A_, and Emax and EC_50_, appear to result from systems with some amplification this concern may not apply. As partial agonists were included in the analysis, an occupancy-response relationship (in a system with some amplification) is likely a close approximation; therefore, the calculations presented herein should be reasonable.

### Potency comparisons and interpretations

It is noteworthy that some partial agonists produced EC_50_ values that are orders of magnitude apart (SS-Table 3, **Figure 2**) although in theory these values should be much closer together as partial agonists occupancy-response curves are closely related. It is possible that ligand binding kinetic properties are leading to overestimates of potency in certain assays; this would also artificially affect the perception of bias ^*24*^. The authors demonstrate that some association kinetics were observed to be delayed but dissociation appears to be rapid, as it was readily reversed by naloxone or DAMGO, suggesting that potency was not greatly exaggerated (SS-Figures S2 and S4) ^*1*^. However, it is particularly interesting that naloxone reversal of mGsi was incomplete (compared to baseline) for all agonists tested, with the exception of SR-17018 which was completely reversed by naloxone (SS-Figure S2B). This would suggest that dissociation kinetics may be delayed and lead to overestimates of potency for the other compounds (including the reference agonist) for the mGsi assay, which was subsequently used for bias calculations and comparison to the therapeutic window. Indeed, if this were the case then it is possible that all bias estimates could be significantly underestimated due to lack of reference agonist equilibrium and overestimation of reference agonist potency.

**Figure 2.**
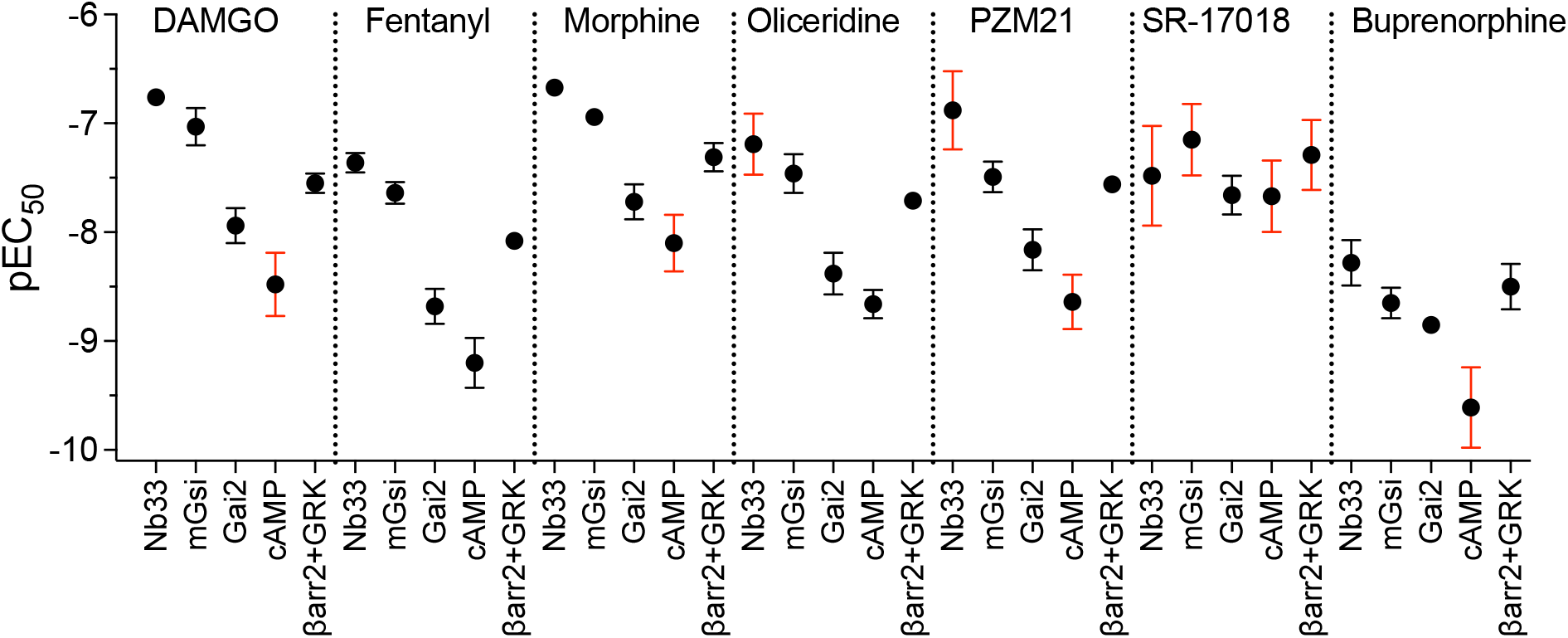
Potency values (pEC_50_) with standard error of the mean are graphically represented from SS-Table 3. The red error bars indicate S.E.M. values that are one half log order or greater. The SS-manuscript states that means are the average of 3-14 experiments.

### Comparison of efficacy (τ values) derived from provided log(τ), Emax and log(τ/K_A_)

For the purposes of bias analysis in the SS-manuscript, the authors focus on five specific responses (Nb33, mGsi, Gαi2 activation, cAMP, and βarrestin2 (+GRK2) recruitment) (see **Table 1** for description of assays). βArrestin2 recruitment is measured in the presence of overexpressed GRK2 as the authors propose that GRK2 overexpression does not alter the analysis of bias (SS-Figure 9). The specific bias analysis that was employed is a reparameterization of the operational model that produces a transduction coefficient, log(τ/K_A_) ^*25, 26*^. When compared to a reference agonist, this efficacy and affinity integration can be used to normalize and consolidate agonist activity into a single composite parameter, namely the normalized transduction coefficient Δlog(τ/K_A_). Further, the analysis used in the SS-manuscript assumes that the responses measured exhibit concentration-response relationships with a Hill slope constrained to one. As such, the authors state that the systems have minimal amplification due to the stochiometric measurement of protein-protein interactions. Constraining the Hill slope to one can be appropriate as log(τ/K_A_) values have been shown to remain linear across a wide range of Hill slope values including one ^*26*^. Of note, upon producing simulated concentration response curves to reflect the data in the SS-manuscript, the Hill slopes do not appear to be broadly variable (**Supplemental Figures 1 and 2**).

When the normalized transduction coefficient was determined in the SS-manuscript, an estimate for agonist affinity (K_A_) is also produced. It can be seen that with Log(τ/K_A_) and K_A_ known, τ and log(τ) can subsequently be determined by Equations 2 and 3:

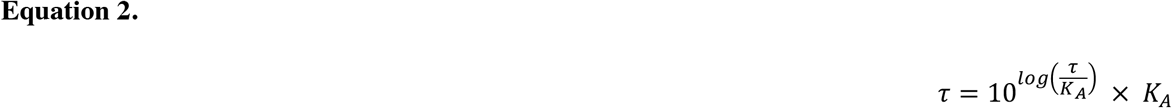

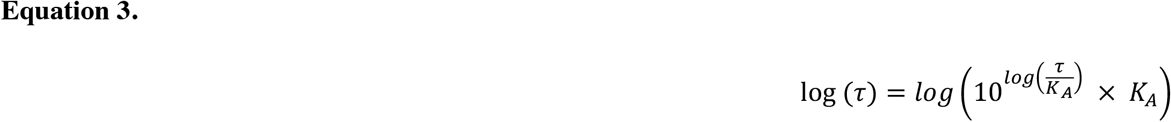

A recent review article highlighted the relationship between the three (or four)-parameter equation and the values that result from the operational analysis ^*20*^. This follows from the parameter definition originally described by Black and Leff, ^*18*^ in Equation 1 (shown above) and Equation 4:

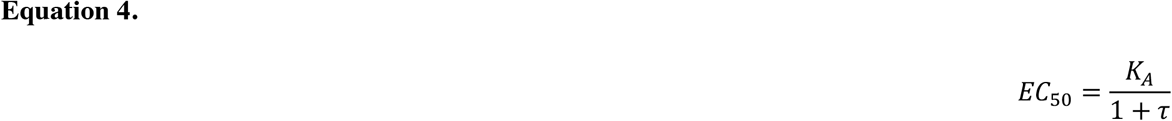

It is possible to estimate K_A_ using the EC_50_ values from SS-Table 3 by applying Equation 5:

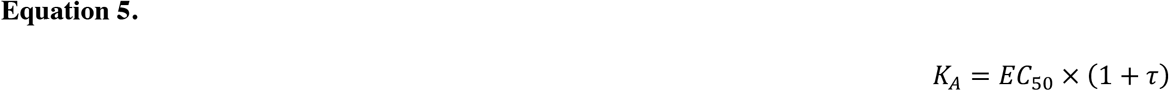

More interestingly, since log(τ/K_A_) was produced for a number of agonists in SS-Table S1, it is possible to solve for τ directly by using Equation 6:

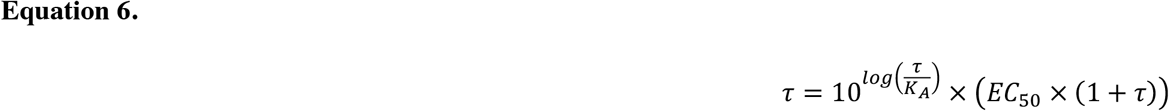

It should be pointed out that these equations are most appropriate for partial agonists as the Emax of a partial agonist is defined at 100% occupancy. Therefore, τ was only estimated for agonists that produced less than 90% efficacy wherein the maximum response could be assumed to be at levels that saturate the available receptor population. Using the log(τ/K_A_) for the partial agonists (SS-Table S1), the Emax (SS-Table 2), and the pEC_50_ (SS-Table 3), the τ parameter can be independently determined; a comparison of these derivations of τ are shown in **Figure 3**. Immediately, it is evident that there is poor agreement between τ values derived from the three sources for the Gαi2 and cAMP measures and this was least consistent for SR-17018.

**Figure 3.**
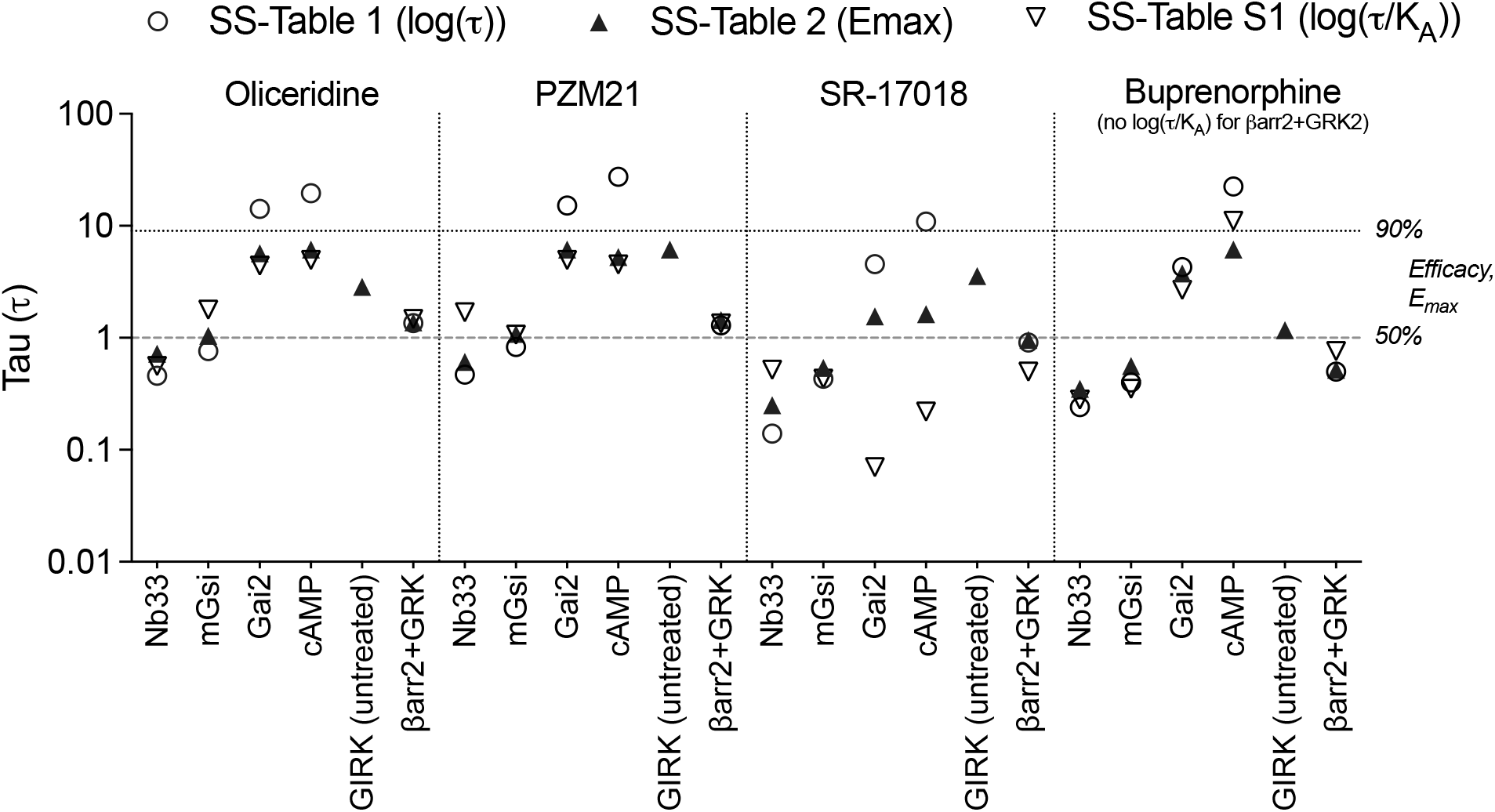
Tau (τ) is graphically presented after derivation from log(τ) (SS-Table 1), Emax (SS-Table 2) or from log(τ/K_A_) (SS-Table S1) using Equations 1–6.

As the focus of the SS-manuscript was the correlation of agonist efficacy or bias with therapeutic window, it was reasonable to produce the Δlog(τ/K_A_) values that are used to evaluate functional selectivity (bias). As such, an estimate of log(τ/K_A_) can be derived from the three-parameter curve fit using Equation 7:

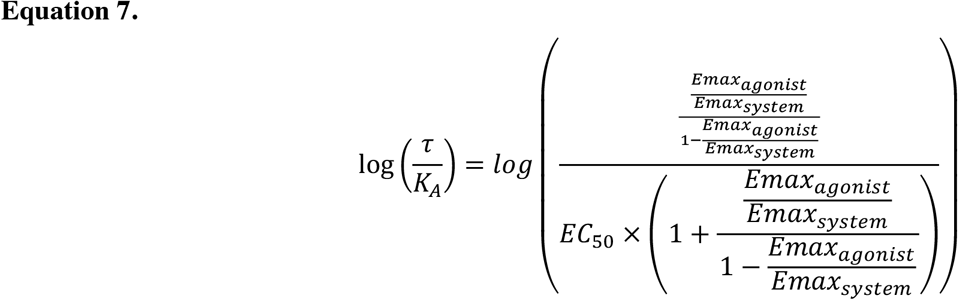

This equation is easily adapted to Microsoft^®^ Excel for both parameter conversion and the further bias analysis that is the principal endpoint. For completeness, all Microsoft^®^ Excel spreadsheets for all figures used for the analyses, herein, are provided for examination in the supplement.

### Comparison of Δlog(τ/K_A_) derived from EC_50_ and Emax compared to Δlog(τ/K_A_) values presented

In bias analysis, the Δlog(τ/K_A_) values specifically provide a discrete measure of each agonists activity in each experimental system. The Δ in the name comes from the correction to the activity of the reference agonist (usually a full agonist where τ cannot be explicitly determined) within each system. The log(τ/K_A_)_Reference_ is defined as the pEC_50_ of the reference agonist in each system and the Δlog(τ/K_A_) for each test agonist is expressed in Equation 8 as:

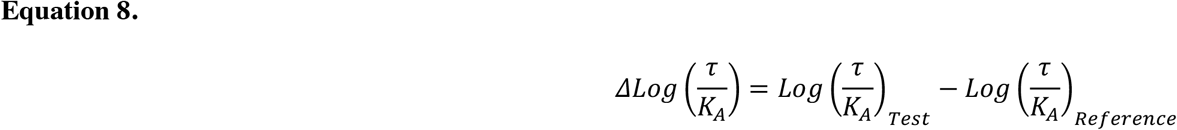

In **Figure 4**, the Δlog(τ/K_A_) for each agonist in each response is plotted on the X-axis (calculated from SS-Table S1). In cases where Δlog(τ/K_A_) was determined for full agonists, the comparison is presented as ΔpEC_50*(Test-Reference)*_. The Y-axis presents the Δlog(τ/K_A_) that was produced using equations 7 and 8 with EC_50_, Emax_system_, and Emax values from SS-Tables 2 and 3. The Emax_system_ is defined as the response produced by DAMGO in each system (100%). The coefficient of determination, (R^2^), when the slope is constrained to one, is also presented in each panel to demonstrate the strength of the agreement between Δlog(τ/K_A_) presented in SS-Table S1 and Δlog(τ/K_A_) derived from the SS-Tables 2 & 3. Notably, the values do not agree for the Gai2 (R^2^= 0.3638) and cAMP (R^2^=0.7182) measures; values for SR-17018 are in disagreement in most assays.

**Figure 4.**
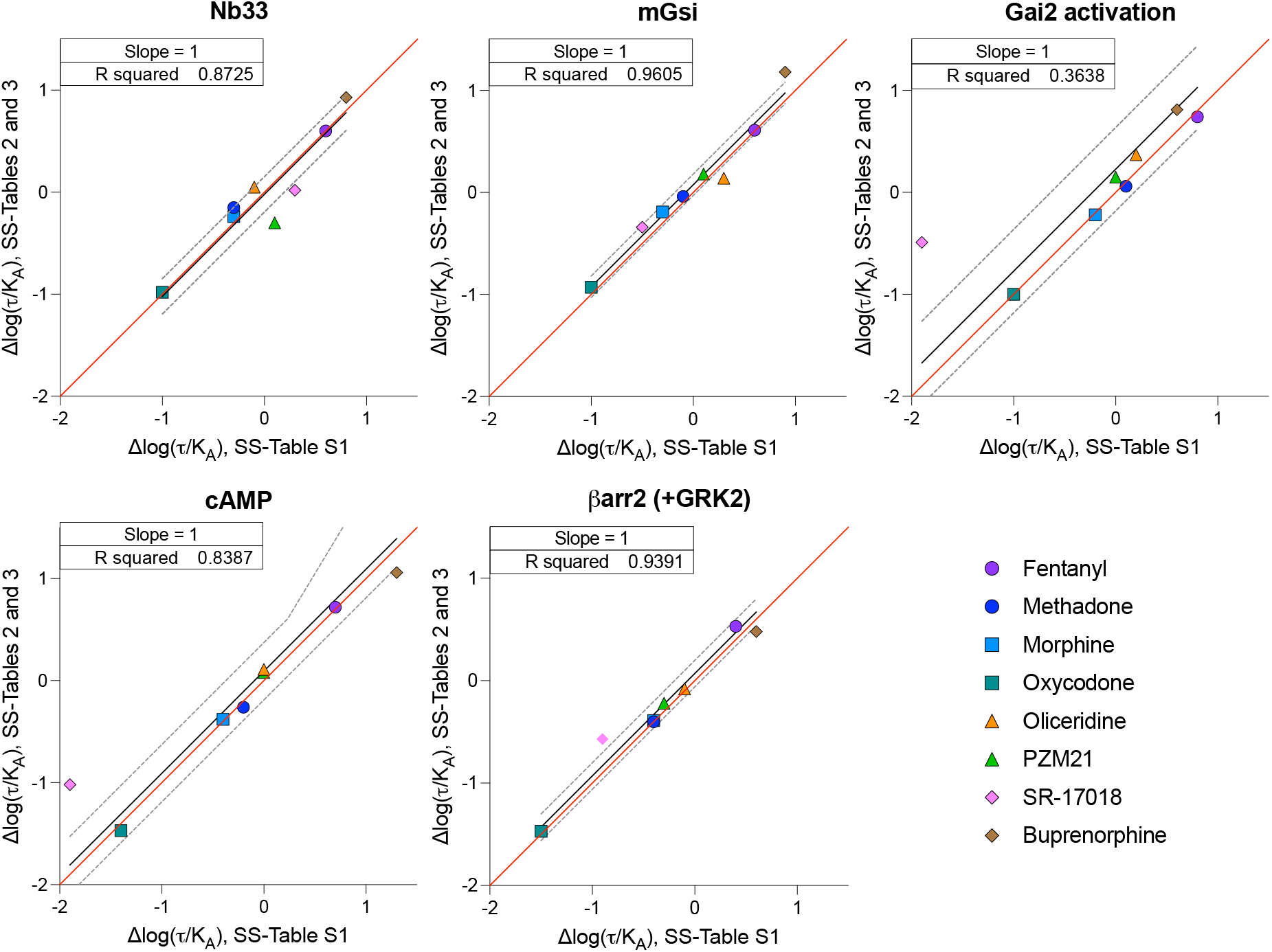
Linear regression of the correlation between the Δlog(τ/K_A_) values derived from potency and efficacy values, presented in SS-Tables 2&3, and the Δlog(τ/K_A_) values derived from the log(τ/K_A_) provided in SS-Table S1. The red line indicates a perfect match of the data points (slope=1), and the black line is the linear fit of the data points with 95% CI shown as dashed gray lines.

### Comparison of ΔΔlog(τ/K_A_) presented in tables, graphs and derived from Emax and EC_50_ values

Having normalized the performance of each agonist in each assay to the performance of the reference agonist DAMGO (mean Δlog(τ/K_A_)), it is further possible to compare an agonist’s relative performance between two assays using Equation 9:

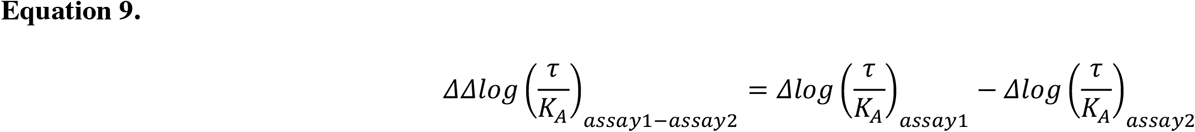

As written in Equation 9, a positive number would indicate a preference for *assay 1* and a negative number would indicate a preference for *assay 2*. When the ΔΔlog(τ/K_A_) and its 95% confidence intervals do not include zero the value is indicative of an agonist that exhibits selectivity between one response and another (i.e. functional selectivity, biased agonism) ^*26*^. More broadly stated, ΔΔlog(τ/K_A_) ≠ 0 indicates that the agonist promotes a discrete selection of an active state that prefers one effector over another; as such ΔΔlog(τ/K_A_) is used to quantify “ biased agonism.” The ΔΔlog(τ/K_A_) is frequently expressed as extending toward or away from one response or another, depending on perspective of preferring or not preferring a receptor active state. This concept was first proposed mechanistically by a number of studies both in brain and recombinant cell systems ^*17, 27*^.

Therefore, using the process described above, the bias factor (as ΔΔlog(τ/K_A_)), can be calculated from the data included in SS-Tables 2, 3 and S1. First, using SS-Table S1, the Δlog(τ/K_A_) value of βarr2+GRK2 can be subtracted from the Δlog(τ/K_A_) for each of the G protein signaling pathways to generate the ΔΔlog(τ/K_A_) (**Equation 9**) to determine preference over βarr2+GRK2 recruitment. These calculated parameters (ΔΔlog(τ/K_A_) are presented in **Figure 5A** and are compared to values graphed in SS-Figure 5 (as estimated from the graphic because the ΔΔlog(τ/K_A_) values were not included in the SS-manuscript). Buprenorphine was not included in this analysis as a log(τ/K_A_) value for buprenorphine in the βarr2 assays was not provided in SS-Table S1. Although the error of these calculations is not readily accessible, the error propagation does not directly inform or affect the means comparison, as discussed above.

**Figure 5.**
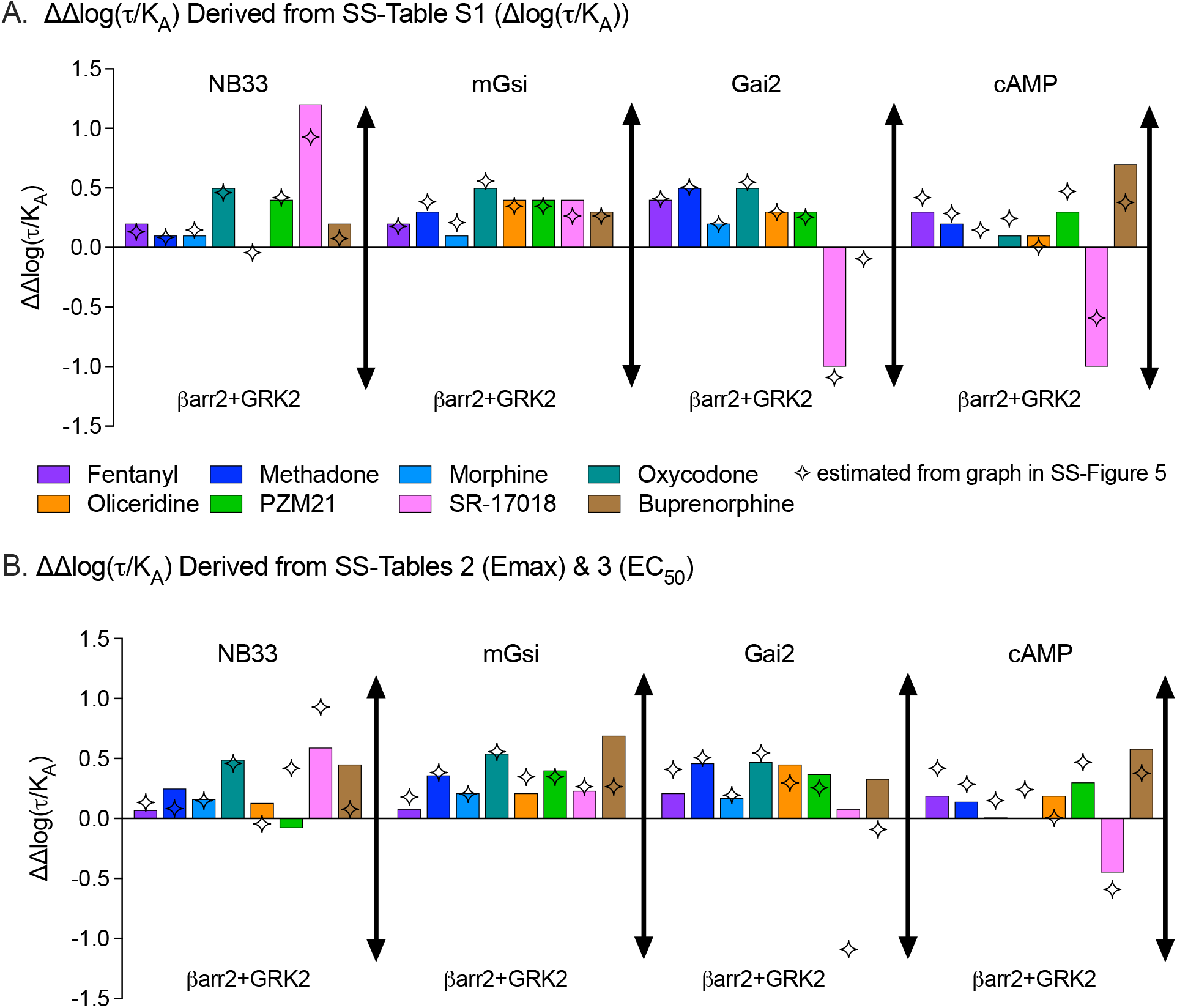
Bias factor analysis presented by calculated ΔΔlog(τ/K_A_) from **A**. SS-Table 1 (Δlog(τ/K_A_)) or **B**. from Δlog(τ/K_A_)) values calculated using the potency and efficacy presented in SS-Tables 2&3. The four-point star represents the values published in SS-Figure 5 as determined by graphical overlay trace estimation. The figure legend is the same for A and B.

Given the discrepancy between calculated ΔΔlog(τ/K_A_) values and those graphed in SS-Figure 5, and a concern that perhaps the SS-Table S1 data were incorrectly derived in the published paper, Δlog(τ/K_A_) values were then calculated from the Emax and EC_50_ values presented in SS-Tables 2 & 3 as described in **Equation 7**. The bias factors (ΔΔlog(τ/K_A_)) were again produced by simple subtraction (**Equations 8 & 9**) and are presented **Figure 5B**. The calculations that produced all values determined in **Figure 5** are provided in the supplemental Microsoft^®^ Excel files.

The comparison of ΔΔlog(τ/K_A_) (**Figure 5**) taken from the three different sources in the manuscript (SS-Figure 5 (simulation), SS-Table S1 (subtraction), and SS-Tables 2&3 (**Equation 7** using Emax and EC_50_), reveal that values are fairly consistent but with some striking exceptions. The values presented in SS-Figure 5 consistently underestimate the G protein signaling preference for SR-17018 across all assays compared to the data obtained from SS-Table S1 (**Figures 5A & B**). The SS-manuscript concludes that SR-17018 “ showed no statistically significant bias toward or away from any G protein activation measure.” If the same error in SS-Figure 5 were applied to the newly calculated ΔΔlog(τ/K_A_), the 95% confidence interval (which spans more than two orders of magnitude) would overlap zero even though the new mean calculated ΔΔlog(τ/K_A_) changes substantially (by an order of magnitude, in some cases). As the 95% confidence intervals for ΔΔlog(τ/K_A_) values are not symmetrical (SS-Figures 5, S6), it is not clearly described how these intervals were determined. However, the standard method proposed^*26*^ was not used, indicating that a direct reassignment to these newly calculated mean values would not be possible or useful. This would lead to the general conclusion that the presences of functional selectivity cannot be proven or disproven, based solely on this analysis, as the error is so large as to preclude *any* meaningful interpretation.

### Comparison of efficacy and therapeutic window

The primary goal of the SS-manuscript was to demonstrate that intrinsic efficacy, and not bias, was responsible for the improvement in the therapeutic window of the biased MOR agonists. To that end, the manuscript produced a series of in vivo experiments to determine therapeutic window as measured by the ΔLogED_50_ between antinociception (hot plate latency) and respiratory suppression (respiratory frequency). These therapeutic window values were then correlated with ligand efficacy, log(τ), as well as functional selectivity (bias, ΔΔlog(τ/K_A_)), between different responses.

For this comparison, the authors focused on intrinsic efficacy in the mGsi assays and bias comparison between mGsi and βarr2+GRK2 assays in SS-Figure 8. **Figure 6** presents a representation of the intrinsic efficacy, log(τ), taken from SS-Table 1 or derived from SS-Table 2 (Emax) graphed for mGsi (**Figure 6A**) and βarr2+GRK2 (**Figure 6B**); this is compared with therapeutic window (**Figure 6C**). It should be noted that it was not possible to derive the log(τ) value for fentanyl in the mGsi assay as the maximum response of fentanyl intersects with the maximal response of the system. As such the log(τ) value for fentanyl, in assays where it approaches or exceeds 100%, will approach infinity. The Emax for the βarr2+GRK2 assay (Emax= 92% in SS-Table 2) was used to estimate log(τ) in **Figure 6B**; no value was provided in SS-Table 1. As the authors point out in SS-Figure 3 of the manuscript, the partial agonists are partial in both G protein signaling assays as well as in βarrestin2 recruitment. It can therefore be interpreted that decreasing the intrinsic efficacy for recruiting βarrestin2 as well as G protein correlates with an improved therapeutic window (**Figure 6A-C**). However, in both cases, there is not much refinement in the assessment, as PZM21 shows a wider therapeutic window, yet the intrinsic efficacy is the same as that observed for oliceridine in both assays.

**Figure 6.**
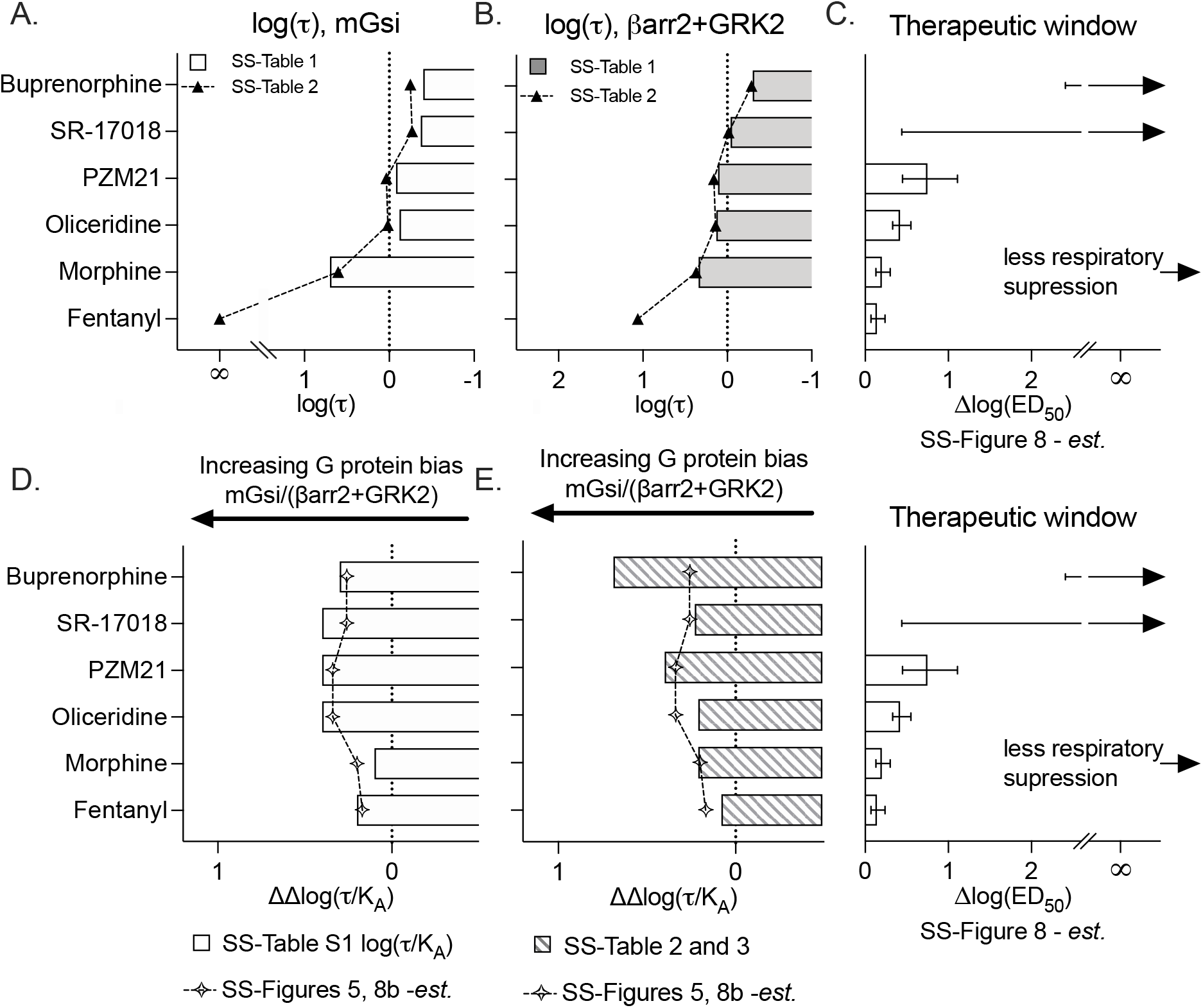
Graphical representation agonist log(τ) and bias factors (ΔΔlog(τ/K_A_) in comparison to therapeutic index as presented in SS-Figure 8. Log(τ) values were plotted from SS-Table 1 or derived from SS-Table 2 and plotted for **A**. mGsi and **B**. βarr2+GRK2 assays and aligned for comparison with the **C**. therapeutic window and error values estimated from SS-Figure 8. Infinity indicates an inability to estimate the mean and error bars are provided as an indication of the lower confidence interval with an arrow pointing towards infinity according to SS-Figure 8 **D**. ΔΔlog(τ/K_A_) values calculated from the Δlog(τ/K_A_) values provided in SS-Table S1 are compared to the values presented in SS-Figures 5 and 8B (estimated). **E**. ΔΔlog(τ/K_A_) values derived from EC_50_ (SS-Table 3) and Emax (SS-Table 2) values as described in the text. The therapeutic window plot is shown again for visual alignment.

Since therapeutic window values were not included in the manuscript, the plot in **Figure 6C** was created by tracing and estimating values that correspond with SS-Figure 8B; error is also included based on this same method. Additionally, the mean therapeutic window values for SR-17108 and buprenorphine were not included as an estimate of ΔED_50_ could not be calculated (as these agonists did not produce 50% of maximum respiratory suppression). Therefore, it is not possible to estimate either the mean or the error for the ΔED_50_ for both SR-17018 or buprenorphine as their means approach infinity. The therapeutic window for these compounds was not measured; any representation of therapeutic windows that does define either of these parameters is, at the minimum, an overinterpretation of the data. With this recognition, the therapeutic window graph in **Figure 6C** presents the lower confidence intervals for means that extend to infinity as depicted in SS-Figure 8.

**Figure 6D and 6E** are from **Figure 5** and are plotted, as presented in SS-Figure 8, for comparison to the therapeutic window plot (shown again within the figure). **Figure 6D** shows the Δlog(τ/K_A_) values calculated from SS-Table S1 (log(τ/K_A_)) and these are shown in comparison to the values that were presented in graphical form in SS-Figures 5 and 8B. Here it can be clearly seen, using the Emax and EC_50_ values to derive log(τ/K_A_), that the biased ligands show an improvement in therapeutic window, moreover, the difference in bias between PZM21 and oliceridine now reflect the differences observed in therapeutic window.

### Comparison of relative intrinsic activity and therapeutic window

In this reanalysis of the data presented in the SS-manuscript, efforts were made to rederive τ and K_A_ from E_max_ and EC_50_, presented in the paper, to arrive at the ΔΔlog(τ/K_A_) bias factors (**Figure 5**). However, given that these are partial agonists, it is also possible to use the values directly produced by the three-parameter concentration-response function to determine relative intrinsic activity ^*25*^ in Equation 10:

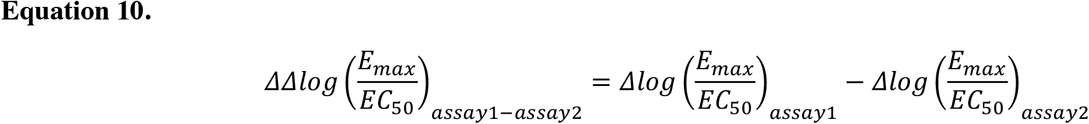

Relative intrinsic activity provides a means of interpreting the activity of agonist in different systems without necessarily fully characterizing the stimulus-response relationship. As such, it is an empirical method of identifying agonists that show preference for a subset of responses compared to other agonists that remain impartial between different responses. The results of this analysis can be visualized in **Figure 7A** comparing ΔΔlog(E_max_/EC_50_), where a preference for G protein signaling over βarr2 (+GRK) is observed across the four platforms used to assess G protein signaling. This analysis is in direct agreement with the ΔΔlog(τ/K_A_) values derived from analysis of E_max_ and EC_50_, to generate τ and K_A_ in **Figure 5B**.

**Figure 7.**
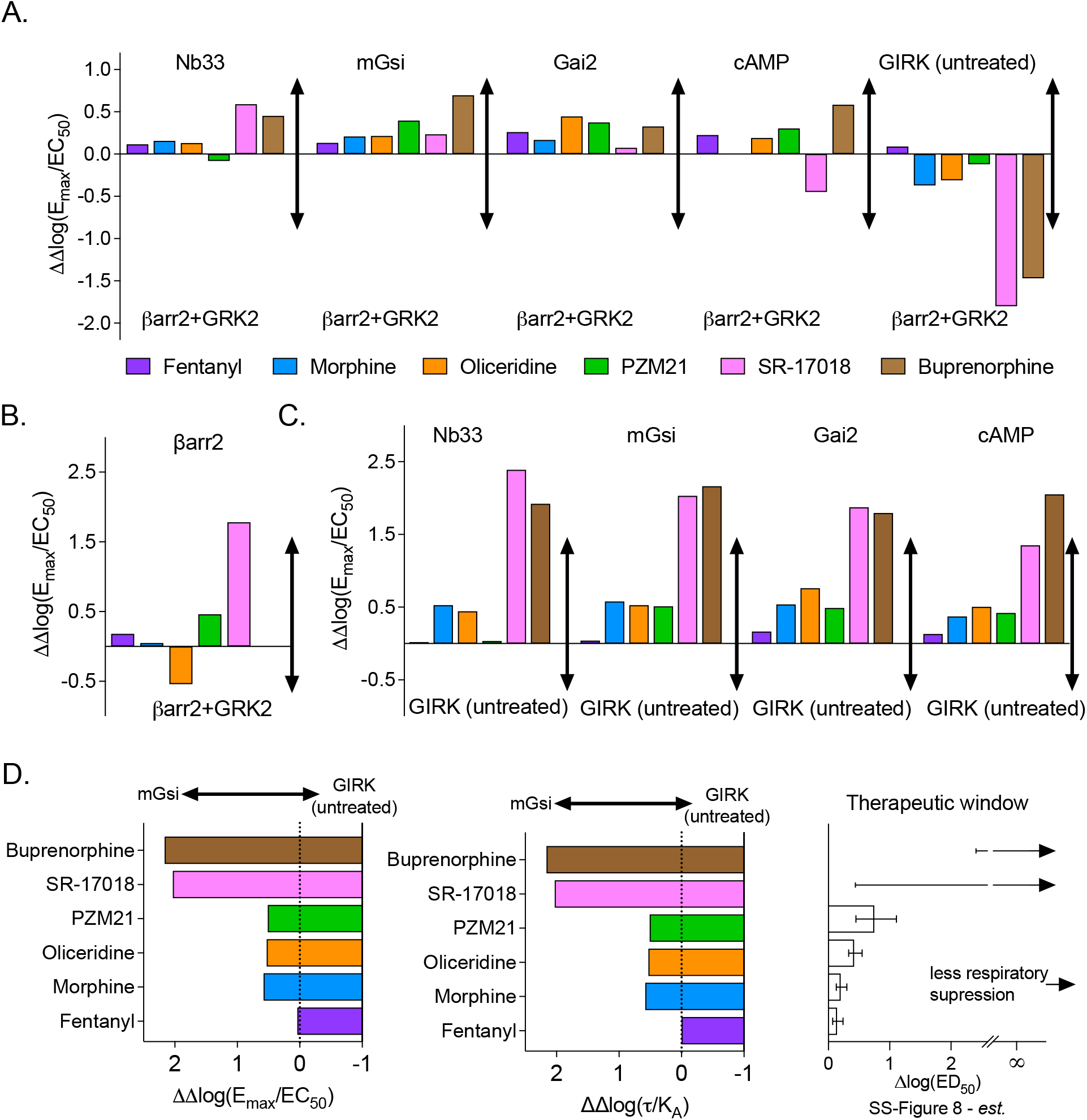
Bias factors determined as ΔΔlog(Emax/EC_50_) from values provided in SS-Tables 2 & 3 comparing responses between **A**. the indicated assays and βarr2 + GRK2 as ΔΔlog(Emax/EC_50_)_*[G protein assay – βarr2+GRK2 assay]*_; **B**. βarr2 vs. βarr2+GRK2 as ΔΔlog(Emax/EC_50_)_*[βarr2 assay – βarr2+GRK2 assay]*_; and **C**. between the indicated assays and GIRK (untreated) as ΔΔlog(Emax/EC_50_)_*[G protein assay – GIRK untreated assay]*_. **D**. Comparison of the as ΔΔlog(Emax/EC_50_)_*[mGsi assay-GIRK assay]*_ and as ΔΔlog(τ/K_A_) _*[mGsi assay-GIRK untreated assay]*_ to the therapeutic window from SS-Figure 8 as also shown in Figure 6.

The authors chose to focus on comparing responses to βarr2 only in the presence of GRK2. GRK2 overexpression is included in the βarr2 assay to improve system sensitivity, as GRK2 phosphorylates MOR and facilitates βarr2 recruitment ^*1, 28*^. Therefore, it is expected that while GRK2 overexpression will enhance agonist potency and efficacy, the degree of enhancement should be *consistent* between the agonists. Preference for βarr2 recruitment in the absence of GRK, compared to GRK overexpression, is presented in **Figure 7B**. Not only is there divergence from unity (ΔΔlog(E_max_/EC_50_) ≠ 0); it is readily apparent that a number of agonists exhibit a preference for βarr2 recruitment in the *absence* of GRK2 relative to their activity in the presence of GRK2 overexpression. This would suggest that GRK2 overexpression could be acting to *decrease* system sensitivity and diminish the βarr2 recruitment produced by the agonist, which is in direct contrast to the expected role of GRK2 in this system (to facilitate βarr2 recruitment). Overall, it is unusual that GRK overexpression worsens an agonist’s ability to recruit βarr2 to the receptor.

We also note that most of the agonists show a preference away from GIRK recruitment and towards βarr2 +GRK2, therefore we asked whether the agonists would demonstrate functional selectivity between the G protein signaling assays. While GIRK was considered an orthogonal assay for G protein-mediated signaling, the data demonstrate robust signaling bias between the ability to activate the other G protein pathways over GIRK (**Figure 7C**). This is particularly intriguing as activation of GIRK has been implicated in MOR-mediated respiratory suppression ^*29*^; indeed, as presented in **Figure 7D**, the bias away from GIRK activation aligns well with the therapeutic window spectra presented in **Figure 6**.

### Active state affinity and rank order efficacy are not preserved

Because the conclusions of the SS manuscript are that intrinsic activity is directly related to therapeutic index, it was relevant to directly compare intrinsic efficacy observed across the different assays. Moreover, since most of the ligands of interest generally act as partial agonists, it was reasonable to generate the active state selectivity predictions that would result from analysis with the two-state model ^*17*^. The active state selectivity ratio (α) can be calculated ^*15*^ as shown in Equation 11:

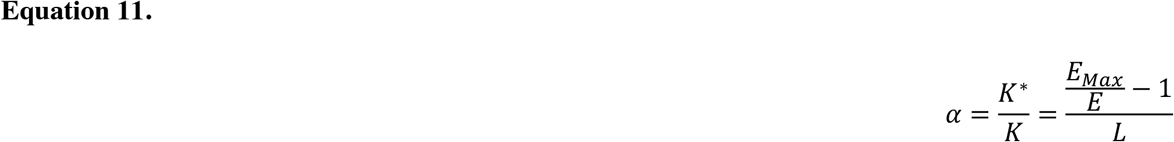

where K* is the active state affinity and K is the inactive state affinity. E_max_ is the maximum response of the full agonist and E is the maximum response of the agonist of interest (from SS-Table 2) and L is the system intrinsic isomerization ratio (R/R*) and was estimated to be 100 for all responses ^*15, 17*^. Further, with the calculated active state selectivity ratio (α), it is possible to use the EC_50_ to calculate the active state affinity constant of the agonist for the receptor as shown in Equation 12 ^*17*^:

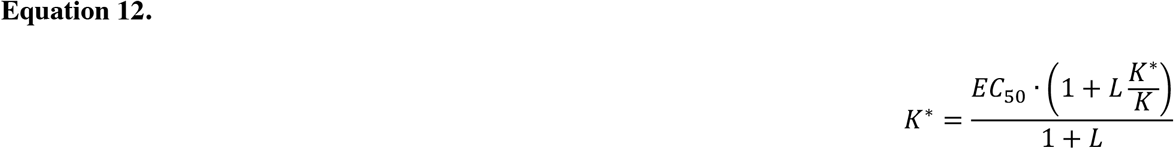

where K* is the agonist affinity constant for the active state in each response using the EC_50_ that is produced from each response. The K*/K is calculated in Equation 11 where, again, L is set equal to 100.

Conceptually, the potency and efficacy of full agonists are highly sensitive to receptor reserve and system sensitivity. That is, a small number of receptors saturated with a full agonist can lead to a full response of the system at a concentration that is less than the active state affinity constant. Partial agonists are less subject to this type of system dependent amplification as their activity is directly proportional to their occupancy. For this reason, it would be expected that the active state affinity (K*) of a partial agonist would be generally conserved across multiple assays, despite differences in signal amplification. However, **Figure 8A** shows that the active state affinity (K*) of several of the partial agonists studied spans two orders of magnitude. This suggests that the differences in affinity may be due to multiple active states of the receptor as previously proposed ^*17*^.

**Figure 8.**
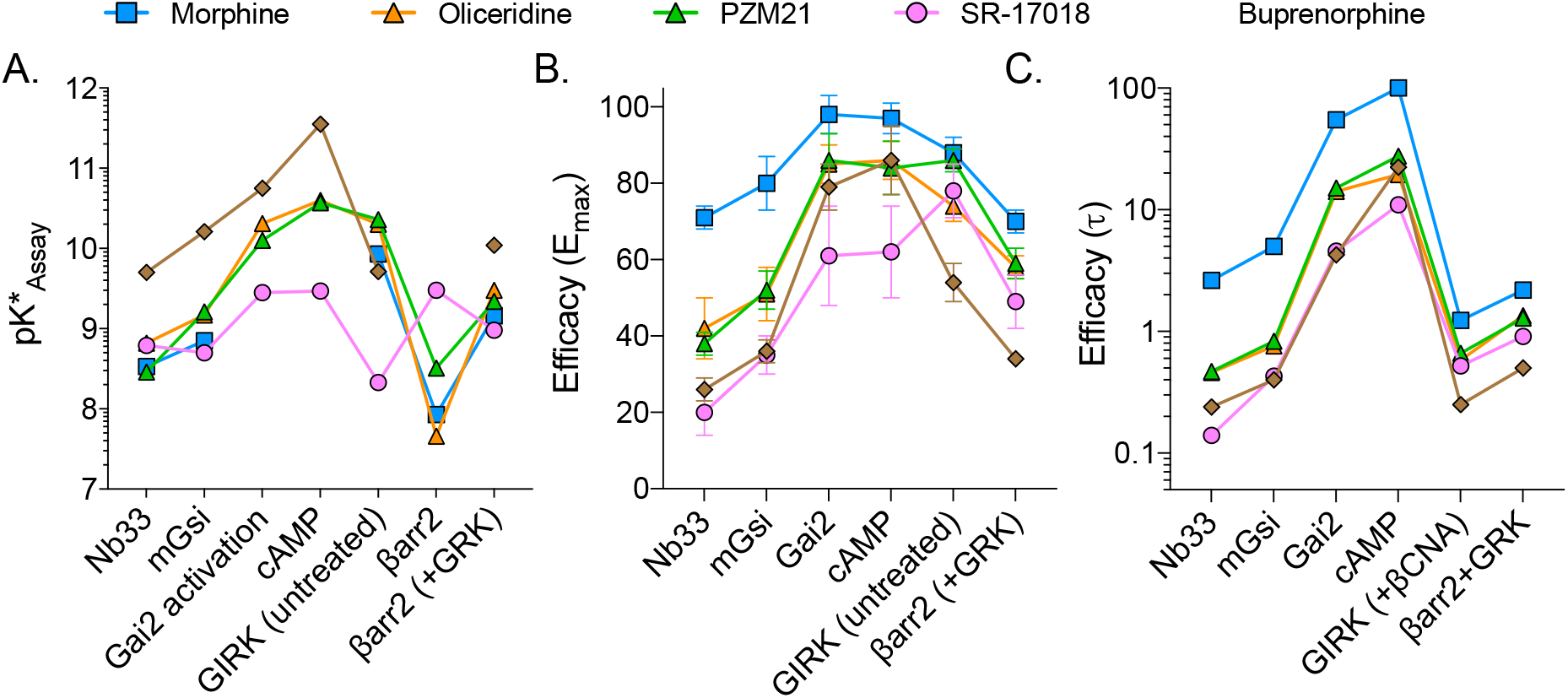
Comparison of intrinsic efficacy of each agonist in the indicated assays as determined by **A**. calculated pK*, (potency values were not included for morphine at Gai and cAMP and buprenorphine at βarr2). **B**. Emax with SEM from SS-Table 2, **C**. τas shown in Figure 2 as derived from log(τ) (SS-Table 1). Missing values in A for morphine are indicated by interrupted lines. Changes in rank order efficacy are easily visualized by the intersection of lines.

However, even if partial agonists could exhibit different affinities for the active state of the receptor (due to system-dependent parameters including receptor-reserve or threshold sensitivity), *the rank order of agonist efficacy should remain constant in different systems if a single active state receptor is responsible for mediating all responses* ^*16, 17*^. **Figures 8B-C** present the comparison of efficacy as E_max_ and τ (derived from SS-Table S1) where it is clear that the relative intrinsic activity (also known as rank order efficacy) is not conserved across the assays. It should be noted that estimates of intrinsic efficacy are not subject to agonist binding kinetics as these values are determined at saturation. These findings distinctly and explicitly implicate more than one active state of the receptor being responsible for stimulating different responses; moreover, the differences in active state selectivity are directly related to the distinction of biased agonists.

## Discussion

The comparison of agonist-induced activity across multiple assays to determine functional selectivity can be a daunting task. While many assays are developed to enhance signal to noise ratios, this can come at a cost of signal amplification. Moreover, certain assays are not amenable to developing stable cells lines and necessitate repeated transient transfections, which can in turn alter receptor number and assay sensitivity. Therefore, it is imperative that comparisons are always made to a reference agonist’s performance (or more than one reference agonist) and that assays are performed in parallel (to avoid changes in the systems over time). The most important requirement for determining functional selectivity is actually two-fold: high quality consistent data and the appropriate application of mechanistic pharmacology analysis. In this evaluation of existing data, we have presented several known approaches for analyzing data to identify a divergence in receptor signaling across assays and have presented an additional analysis adapted from the two-state model of receptor activation (K*). The benefit of determining K* is that is produces a microaffinity constant for the active state preference over the inactive state of the receptor. This allows for normalization of efficacy and potency to produce affinity estimates. If there is no functional selectivity, it would be expected that the rank order of these affinity estimates would be conserved across all systems that are being investigated. Divergence from this conservation will indicate a preference for an assay-dependent active state. Further, the utilization of mechanistic analysis to underpin more empirical analysis can lend a degree of confidence to the general conclusion that ligand bias is observed within a set of data.

In the re-calculation of bias from the SS-manuscript it was not possible to interpret error for the data set as there was only limited information provided by the authors; for example, the number of experiments were provided as a range (3-14 replications) and the S.E.M. was presented in the tables. However, since the derivation of τ and K_A_ is dependent on the potency (EC_50_) and efficacy (Emax) within each assay the ability to produce a “ bias factor” with any reasonable error of certainty will require that the individual assays are: first, reproducible, and secondly, adequately powered by replicates. There is reasonable concern that the experimental measures presented herein exhibit large error in the estimation of potency (pEC_50_ in SS-Table 3) and efficacy (Emax, SS-Table 2). This can be visualized in **Figure 2**, where the error in potency exceeds a half of log order in several assays. Since the authors report running 3-14 experiments per data set, it would seem that perhaps, in cases where error is very high, that the studies were underpowered. This initial error propagates throughout additional analyses, leading to 95% CI that are greater than the mean (SS-Figure 5, SS-Figure S6). Therefore, it is apparent that several assays reported in the SS-manuscript are underpowered and do not support the conclusion that there is no “ significant” bias observed for any of the compounds. This concern with reproducibility is further supported by the degree of error present upon averaging the parameters derived from DAMGO response curves. DAMGO, as the reference agonist, would be included in every experiment (presumably accounting for the upper n=14 described in SS-Tables 1-3); yet, the margin of error in DAMGO’s potency was wider than expected for several of the signaling assays **(Figure 2**, SS-Table 3). The reproducibility issues may stem from the use of transient transfection systems with variable receptor density and effector expression levels. Moreover, there was no indication that protein and receptor expression levels were monitored for consistency between experiments.

In many ways, the SS-manuscript reproduces observations from several investigators that have proposed that improving G protein signaling while avoiding βarrestin2 recruitment may be a useful approach to limit respiratory suppression while preserving antinociception ^*3-6, 8, 11*^. Specifically, the authors reproduce in vivo observations originally published for SR-17018 ^*8*^, oliceridine ^*11*^, and PZM21 ^*6*^. Moreover, the demonstration that PZM21, at doses up to 100 mg/kg, produces less than half of the respiratory suppression induced by 10 mg/kg morphine; this is very encouraging as it had only been tested up to 40 mg/kg in the first study, where it showed no respiratory suppression compared to vehicle ^*6*^. It should be noted that these observations are in direct contrast to a study that showed PZM21 and morphine producing equivalent respiratory suppression at 10 mg/kg ^*30*^.

The idea that decreasing intrinsic efficacy may serve to improve the therapeutic utility of mu opioid receptor agonist to manage pain and limit side effects has been pursued for some time, resulting in compounds such as buprenorphine ^*31*^. However, caution should be maintained in translating efficacy observed in cultured systems to expectations observed in clinical use. In particular, buprenorphine is fully efficacious in the treatment of some pain conditions ^*32*^, as is morphine, even though both perform as partial agonists across many assay systems, as demonstrated in the SS-manuscript. As such, there has not been a direct correlation between partial agonism observed in different cell-based assays and the effects of a drug *in vivo*; therefore, it may not be fully predictive that a partial agonist may be safer regarding respiratory suppression. Indeed, buprenorphine produces respiratory suppression in humans ^*33*^, with the additional complication that it is difficult to reverse with naloxone ^*34*^ as buprenorphine is a long lasting, high affinity agonist ^*31*^. In rats, buprenorphine produces respiratory suppression to the same extent as fentanyl which would also seem to contradict the efficacy hypothesis as well as the biased agonism hypothesis ^*35*^. However, studies with buprenorphine are further complicated by the fact that it has activity at other opioid receptors and also has nonselective active metabolites that can suppress respiration ^*36-38*^.

The conclusion of the SS-manuscript is that partial agonism is responsible for the avoidance of respiratory suppression, and this is supported by the choice of agonists and how those agonists performed in the G protein signaling assays selected. However, the same could be said for the efficacy in the βarrestin2 recruitment assays, as the biased agonists act as partial agonists there as well, in a manner that also aligns with an improved therapeutic window. The data presented here may support the role of agonist efficacy in determining the extent of respiratory suppression in mice; however, no counter argument was purposefully explored (i.e. testing of a full agonist with less respiratory suppression or a partial agonist with more respiratory suppression).

The study by Schmid et al., 2017, addressed this question by generating a series of structurally related MOR agonists that spanned a spectrum of both bias and intrinsic efficacy ^*8*^. One example from that study is SR-11501, a partial agonist that has similar potency and efficacy to morphine in G protein signaling and in antinociception studies. However, SR-11501 displays bias towards βarrestin2 recruitment over G protein signaling and produces fentanyl-like respiratory suppression, resulting in a very narrow therapeutic window. Moreover, examples of two full agonists that display bias towards G protein signaling, SR-14968 and SR-14969, were shown to produce a wider therapeutic window than morphine (less respiratory suppression vs. antinociception) ^*8*^. Schmid et al. also showed that fentanyl, which has a narrow therapeutic window, acts as a partial agonist in mouse brainstem membranes using ^35^S-GTPγS binding assays ^*8*^. Fentanyl has also been shown to act as a partial agonist in spinal cord ^*39*^ and in rat thalamus ^*40*^ in studies assessing ^35^S-GTPγS binding. Therefore, intrinsic efficacy can be uncoupled from biased agonism and, within the SR series of agonists, G protein signaling bias was shown to correlate with decreased respiratory suppression, independent of intrinsic efficacy ^*8*^.

Regardless of the distinction of bias, the comparison of active state affinity constants and the observation that rank order efficacy was not conserved across the spectrum of assays investigated provides compelling support for the idea that these experimental compounds present interesting and substantially novel pharmacology. Ultimately, the determination of biased agonism, or intrinsic efficacy for that matter, will be dependent on the cellular signaling systems used (i.e. receptor density, effector expression, amplification of signal) as well as reference agonist used to determine the maximum possible signaling response in the system. It is not unexpected that perceptions of bias and efficacy will vary between studies ^*22, 41*^. In a recent example, similar BRET-based studies were elegantly employed for similar analysis by Ehrlich et al., ^*42*^ investigating ligand bias at the mu opioid receptor where they demonstrate diverse signaling profiles of oliceridine, PZM21 and buprenorphine in neurons. In time, the value of these cellular readouts as predictors of agonist function may be supported by detecting state-dependent structural conformations of MOR induced by biased ligands binding to the receptor ^*43-45*^. This does not make their pharmacology less remarkable not does it disprove the hypothesis that ligands may demonstrate active state selectivity. On the contrary, as more studies present interesting and novel findings regarding the spectrum of activity these compounds demonstrate, it may be possible to more definitively establish these differences.

In particular, the demonstration of changes in rank order efficacy is a compelling finding that further substantiates and validates the value of these compounds. From a practical standpoint, however, the correlation between the pharmacological properties and the improvement in therapeutic efficacy while limiting side effects in human studies should drive further investigations. Indeed, oliceridine, which is a partial agonist in both G protein signaling and βarrestin2 recruitment, as well as a biased agonist for G protein signaling, is producing superior analgesia/respiratory suppression in post-operative pain ^*11, 13, 14*^. Therefore, while partial agonism could be a desirable pharmacological property, biased agonism may serve as a means to further improve the therapeutic window.

