## Supplementary material for "Low intrinsic efficacy alone cannot explain the improved side effect profiles of new opioid agonists": Supp Figure 1

Supplemental Figure 1. Simulation of curves from $EC_{50}$ and Emax values.

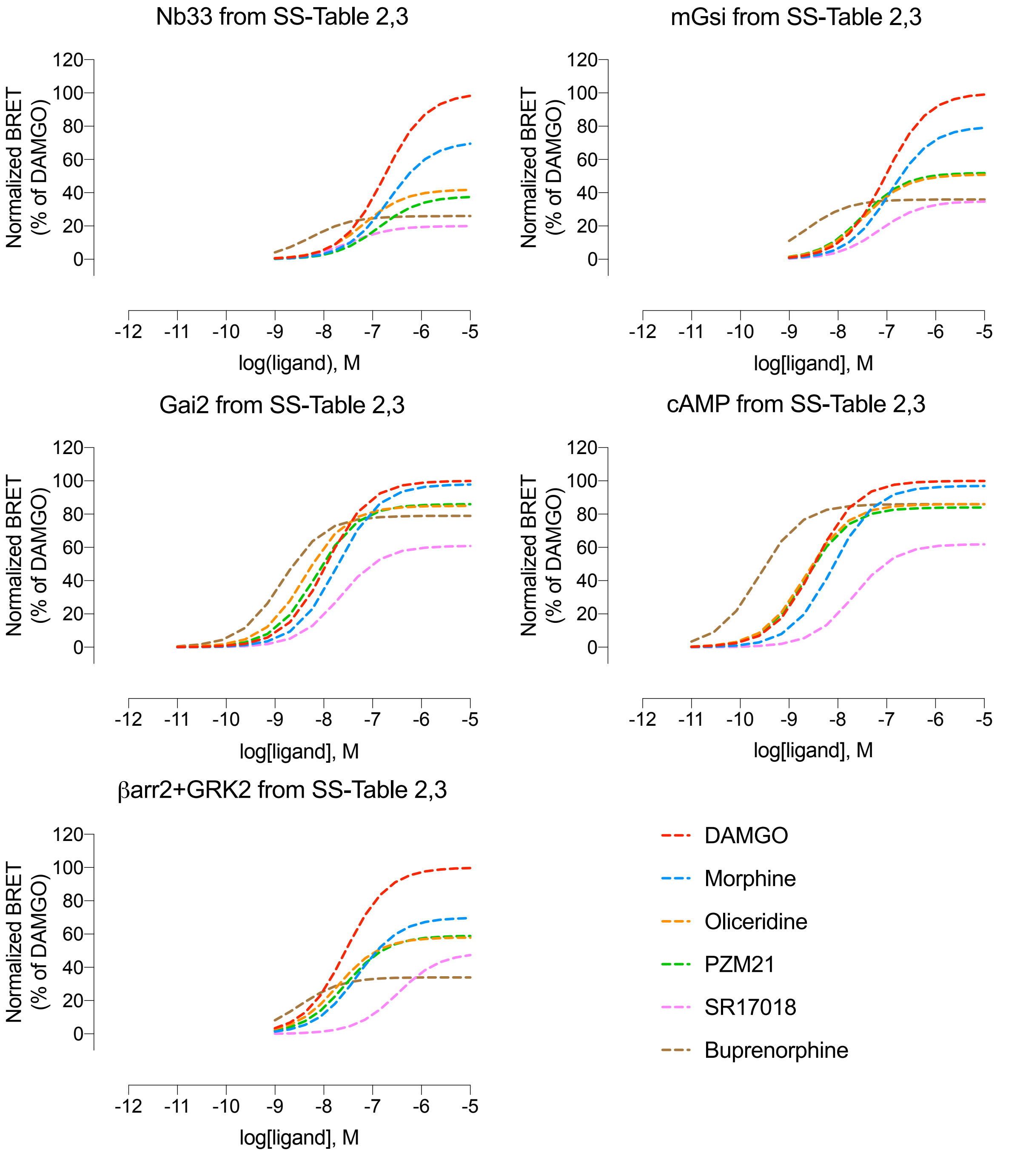


**Supplemental Figure 1.** Simulated concentration response curves generated from SS-Table 2 and SS-Table 3 parameters using 3 parameter nonlinear regression, GraphPad Prism 9.
